# A combined computational-biochemical approach offers an accelerated path to membrane protein solubilization

**DOI:** 10.1101/2023.06.18.545468

**Authors:** Mariah R. Pierce, Jingjing Ji, Sadie X. Novak, Michelle A. Sieburg, Shikha Nangia, James L. Hougland

## Abstract

Membrane proteins are difficult to isolate and purify due to their dependence on the surrounding lipid membrane for structural stability. Detergents are often used to solubilize these proteins, with this approach requiring a careful balance between protein solubilization and denaturation. Determining which detergent is most appropriate for a given protein has largely been done empirically through screening, which requires large amounts of membrane protein and associated resources. Here we describe an alternative to conventional detergent screening using a computational modeling approach to identify the most likely candidate detergents for solubilizing a protein of interest. We demonstrate our approach using ghrelin *O*-acyltransferase (GOAT), a member of the membrane-bound *O*-acyltransferase (MBOAT) family of integral membrane enzymes that has not been solubilized or purified in active form. A computationally derived GOAT structural model provides the only structural information required for this approach. Using computational analysis of detergent ability to penetrate phospholipid bilayers and stabilize the GOAT structure, a panel of common detergents were rank ordered for their proposed ability to solubilize GOAT. Independently, we biologically screened these detergents for their solubilization of fluorescently tagged GOAT constructs. We found computational prediction of protein structural stabilization was the better predictor of detergent solubilization ability, but neither approach was effective for predicting detergents that would support GOAT enzymatic function. The current rapid expansion of membrane protein computational models lacking experimental structural information and our computational detergent screening approach can greatly improve the efficiency of membrane protein detergent solubilization supporting downstream functional and structural studies.

## Introduction

Integral membrane proteins such transporter, receptors, and membrane bound enzymes are essential for biological function.^1^ These proteins are important for importing glucose to the brain, hormone signaling by GCPRs, and lipid modification, to name merely a few of their plethora of roles.^2–4^ The dependence of these proteins on the surrounding membrane lipids for their structural stability and functional activity renders experimental investigations extremely challenging.^5^ A variety of approaches have been employed to isolate integral membrane proteins from cellular membranes, including detergents, cell microsomes, lipid-like polymers, peptides, and lipid nanodiscs.^6^

The most commonly used approach is detergent solubilization, wherein the membrane protein is extracted from cellular phospholipid bilayers by incubation with a detergent at concentrations usually above the critical micelle concentration (CMC).^7–8^ The vast majority of proteins can be solubilized by detergents, as clearly demonstrated by the widespread use of strongly denaturing detergents such as sodium dodecyl sulfate (SDS) for preparing proteins for polyacrylamide gel electrophoresis.^9^ The challenge with membrane protein detergent solubilization is replacing the native membrane lipids with detergent molecules while maintaining the native protein structure. Among the three chemical classes of detergents (ionic, zwitterionic, and nonionic),^10^ nonionic detergents are most popular for membrane protein solubilization as to their lack of charge results in a lower propensity to denature structured proteins. Common examples of “mild” nonionic detergents include Triton and Tween detergents, dodecyl maltoside (DDM), and digitonin.^11^ However, there are examples of membrane proteins that have been solubilized by harsher (more denaturing) ionic detergents like Fos-cholines (FOS) or zwitterionic detergents.^10–11^ Identifying detergents that can effectively solubilize a given membrane protein without protein denaturation remains predominantly an empirical process. Recent advances in high-throughput methods for optimizing membrane protein expression and detergent solubilization have accelerated this process for some targets,^12–14^ but the infrastructure required for these approaches limits their use outside major structural biology centers and requires large amounts of membrane protein for successful screening.

Advances in computational methods for prediction of protein structure have created new avenue for structural insights into uncharacterized membrane proteins. Established homology modeling approaches require a solved structure of a reasonably related protein.^15–16^ Application of metagenomics and coevolutionary contact analysis provide additional pairwise distance constraints for structural modeling.^17–19^ Most recently, machine learning and artificial intelligence-based tools such as Alphafold and its progeny have led to an explosion in predicted protein structures covering many organismal proteomes.^20^ While these modeling approaches offer important information regarding the structures of unsolved proteins, they cannot offer the resolution of experimentally determined protein structures. Similarly, Alphafold and related computational approaches currently offer at best a minimal ability to determine binding contacts between proteins and their ligands, inhibitors, and/or substrates.^21^ To achieve these goals requires solubilization and purification of membrane proteins for co-crystal structures or bound complex analysis by cryo-EM.

In this work, we leverage advances in protein structural modeling combined with computational analysis of detergent-membrane and detergent-protein interactions to create a facile process to identify detergents likely capable of solubilizing an integral membrane protein. We then biochemically validate the output of the computational predictions using an integral membrane protein that has not been successfully solubilized in active form. Our candidate protein is ghrelin *O*-acyltransferase (GOAT), an integral membrane enzyme member of the membrane-bound *O*-acyltransferase family.^22–23^ GOAT octanoylates the peptide hormone ghrelin,^24–25^ which is implicated in appetite stimulation, growth hormone and insulin secretion, metabolic response to starvation, and glucose homeostasis amongst other physiological roles.^26–31^ GOAT has not been solubilized in active form, with solubilization in FOS-16 leading to loss of ghrelin acylation activity.^32–33^ The topology of mouse GOAT was determined by selective permeabilization,^34^ with the same group subsequently reporting an extensive detergent screen without successful solubilization of active GOAT.^32^ Through application of coevolutionary contact analysis coupled with molecular dynamics and biochemical validation, Campaña and co-workers published a computational model of human GOAT (hGOAT) in 2019. This model matched the published mouse GOAT topology, and revealed the helical cluster “MBOAT core” and transmembrane channel now considered a hallmark of MBOAT-family enzymes that modify protein substrates.^23, 35–39^ Despite the lack of hGOAT solubilization and purification, extensive functional studies have determined the substrate selectivity of GOAT and provided insight into the enzyme’s catalytic mechanism.^32, 40–45^ Building on this foundation, solubilization and purification of hGOAT will facilitate functional and structural studies of this enzyme and its acylation activity.

## Results

### Computational screening of hGOAT detergent solubilization

For computational modeling and simulation, hGOAT detergent solubilization was divided into two discrete processes. The first is detergent insertion/invasion of the phospholipid bilayer which is required for the detergent to displace and replace the membrane lipids that directly contact the exterior surface of hGOAT. Performance in this step reflects a detergent’s ability to extract hGOAT from its native membrane context. The second process is detergent stabilization of the folded hGOAT structure to avoid protein unfolding leading to enzyme inactivation. Performance in this step correlates with a detergent’s ability to support the native hGOAT fold required for enzyme activity. For these computational studies, we chose a panel of nonionic and zwitterionic detergents as these two classes are well represented in the structural biology literature (Table 1).

**Table 1.** Detergents used for computational studies and hGOAT solubilization trials

| Detergent | Full Name | Structure |
| --- | --- | --- |
| BOG | n-Octyl-β-D-glucopyranoside | |
| CHAPS | 3-((3-Cholamidopropyl)dimethylammonio)-1-propanesulfonate |  |
| DDM | n-Dodecyl-β-D-maltoside | |
| FOS-12 | Fos choline-12 |  |
| FOS-16 | Fos choline-16 |  |
| GDN | Glyco-diosgenin |  |
| LMNG | Lauryl maltose neopentyl glycol |  |
| MEGA-9 | Nonanoyl-N-methylglucamide |  |

To model detergent invasion of the phospholipid bilayer (Figure 1A), a bilayer of 40 DOPC (dioleoylphosphatidylcholine) and 40 DPPC (dipalmitoylphosphatidylcholine) molecules was built with the CHARMM-GUI Membrane Builder.^46–47^ For each detergent, 240 detergent molecules were inserted with the bilayer into a 12 nm cubic simulation box. From this initial state, the system was energy minimized and equilibrated at room temperature for 1 ns followed by a 500 ns production run. Following this simulation, detergent invasion was determined by measuring the percentage of total detergent molecules in the system that are within 6Å of a membrane phospholipid (Figure 1B). This analysis separated the eight detergents into three groups, with DDM, MEGA-9, and BOG exhibiting the most efficient membrane invasion (>45%). LMNG and FOS-16 were less efficient at penetrating the DOPC/DPPC bilayer (20-45 %), with FOS-12, GDN, and CHAPS showing the lowest percentage (<20%) of detergent molecules in contact with membrane lipids (Figure 1C). We note the most effective detergents are nonionic simple alkylated glycosides; inclusion of charge or more complex glycoside structures reduces the membrane invasion ability.

**Figure 1.**
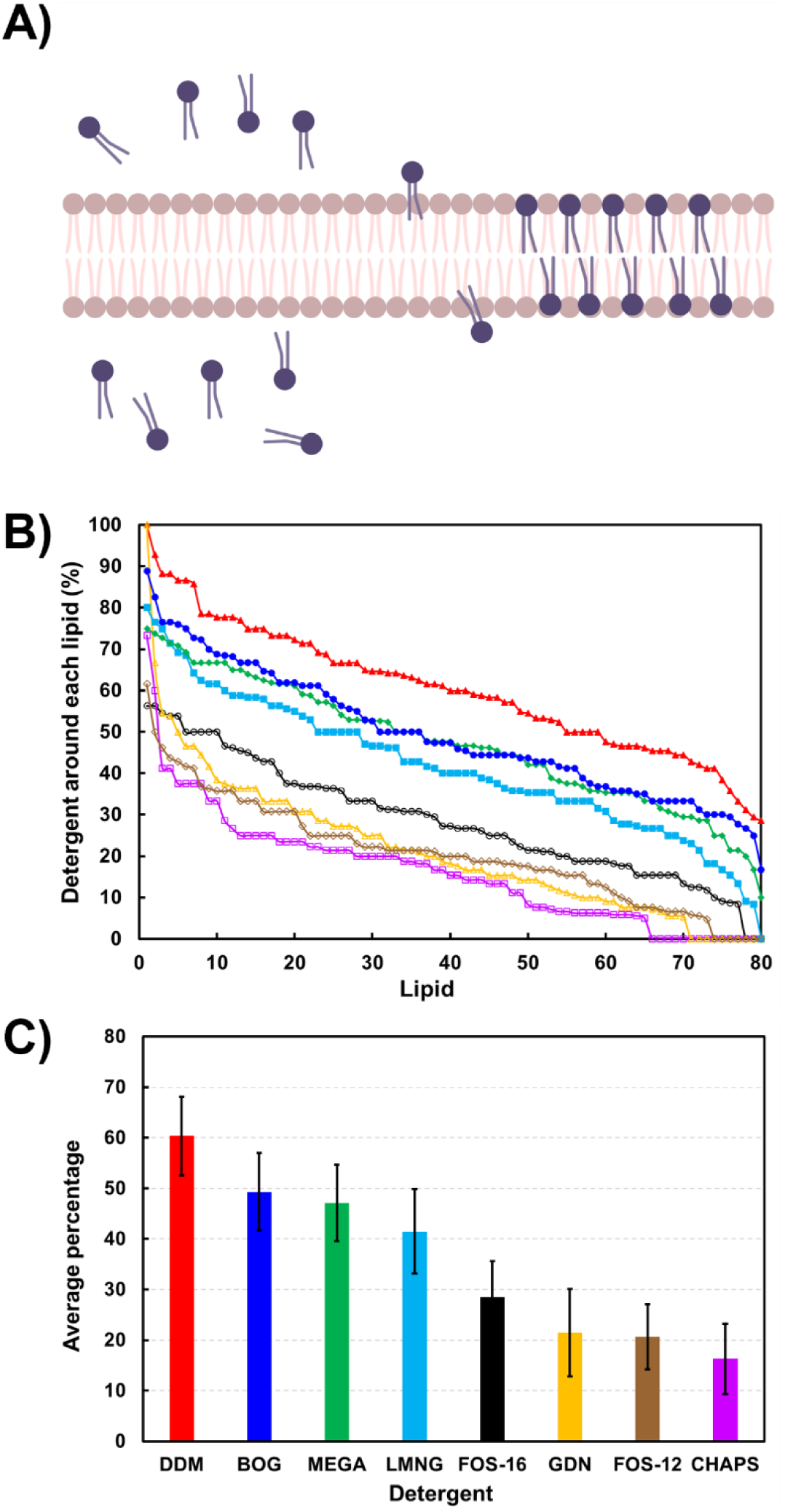
Computational modeling of phospholipid bilayer invasion by detergents. A) Schematic of detergent invasion of DOPC/DPPC phospholipid (light brown) bilayer by solubilizing detergent (blue). Panel made with Biorender.com. B) Percent detergent around each lipid, ranked from highest to lowest population. DDM (red, filled triangle), BOG (blue, filled circle), MEGA-9 (green, filled diamond), LMNG (cyan, filled square), FOS-16 (black, open circle), GDN (yellow, open triangle), FOS-12 (brown, open diamond), and CHAPS (pink, open square). **C)** Average percentage of detergent around each lipid, with same color scheme as panel B. Error bars represent one standard deviation.

To study the stability of the hGOAT structure with detergents, another eight simulation systems with a box length of 15 nm were built, each containing one GOAT protein and 490 detergent molecules. The generation of the hGOAT structure computational model was described in our previous work.^17^ Following energy minimization and equilibration, a 200 ns atomistic molecular dynamics run was performed, and protein stability was determined by the average root mean square fluctuation (RMSF) of hGOAT during the simulation (Figure 2A). Analysis of average ΔRMSF GOAT structural stabilization by detergents relative to membrane-equilibrated structure yielded a distinct rank for the eight examined, with FOS-12, CHAPS, and LMNG supporting the lowest ΔRMSF (Figure 2B). MEGA-9 and FOS-16 yielded comparable ΔRMSF values higher than FOS-12, CHAPS, and LMNG. Finally, the last three detergents (GDN > DDM > BOG) led to higher ΔRMSF values consistent with the destabilization of the hGOAT folded structure.

**Figure 2.**
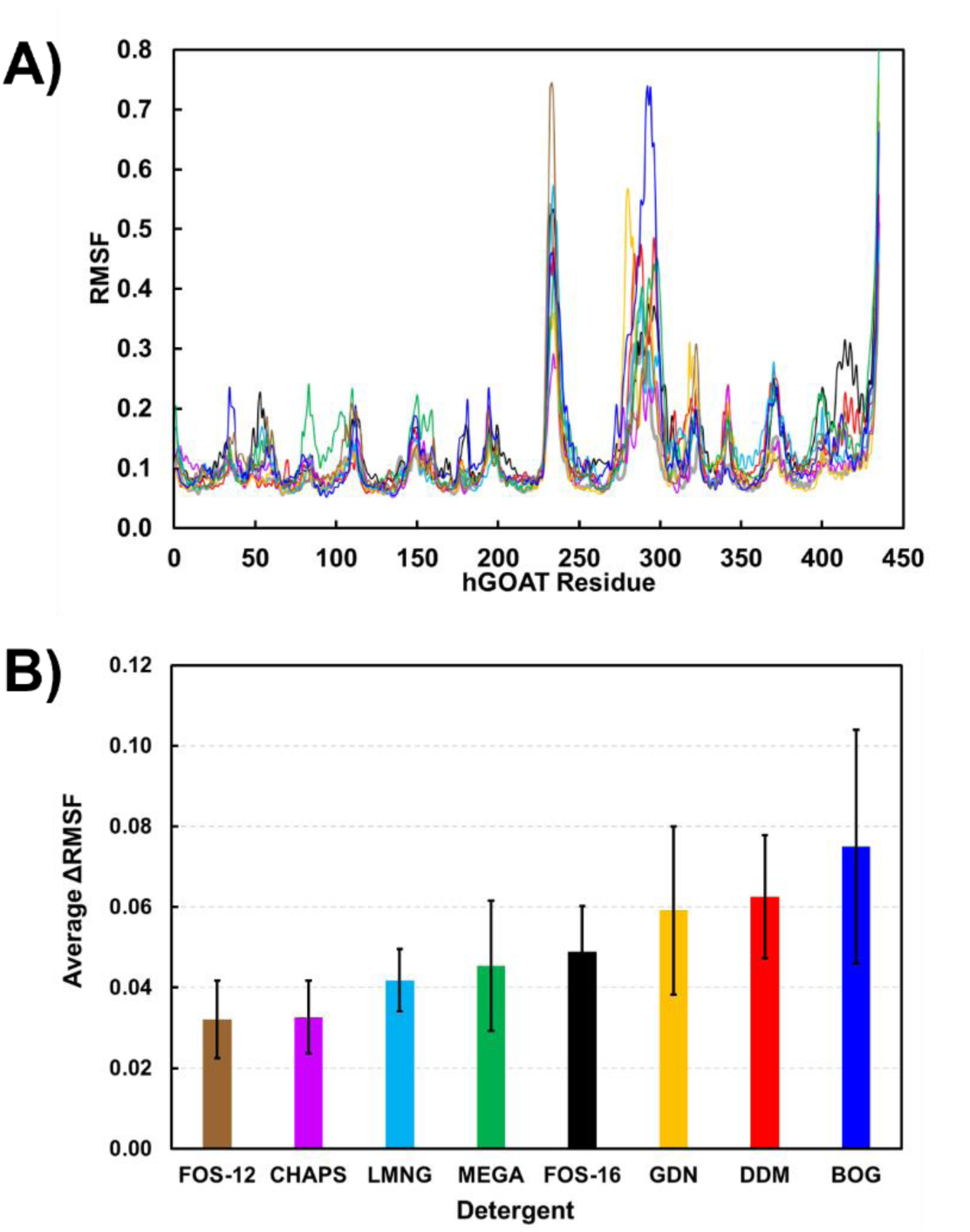
hGOAT structure stabilization by detergents. **A)** Root mean square fluctuation (RMSF) of hGOAT residues. **B)** average **Δ**RMSF for each detergent. DDM (red), BOG (blue), MEGA-9 (green), LMNG (cyan), FOS-16 (black), GDN (yellow), FOS-12 (brown), and CHAPS (pink).

Combining the membrane invasion and hGOAT stabilization data reveals that MEGA-9 and FOS-16 are effective membrane solubilizers because they both penetrate the membrane and stabilize the structure of the extracted protein. Other detergents are either strong invaders and poor stabilizers or vice versa (Figures 1C and 2B). The result with FOS-16 is particularly interesting, as this detergent was reported to efficiently solubilize mouse GOAT but did not support enzymatic activity.^32^ This finding supports the value of computational studies to potentially identify detergents that can support both membrane protein solubilization and maintenance of the native protein fold required for biological function.

To further explore this issue, we examined the structural alignment of hGOAT from a membrane-embedded simulation with FOS-16 and MEGA-9 and detergent-stabilized structures (Figure 3). In both detergents, the structures deviate from the native membrane protein structure, but FOS-16 causes structural changes in the loops and transmembrane (TM 11) regions, whereas MEGA-9 affects the cytoplasmic loop (Figures 3A-B). Further investigation of detergent-hGOAT interactions revealed that these two detergents exhibit distinct regioselectivity in their contacts with the enzyme (Figures 3C-D). Our analysis showed FOS-16 primarily interacts with the hGOAT transmembrane helical regions via its hydrophobic alkyl chain. In contrast, MEGA-9 interactions with hGOAT involve its hydrophilic head and hydrophobic tail groups.

**Figure 3.**
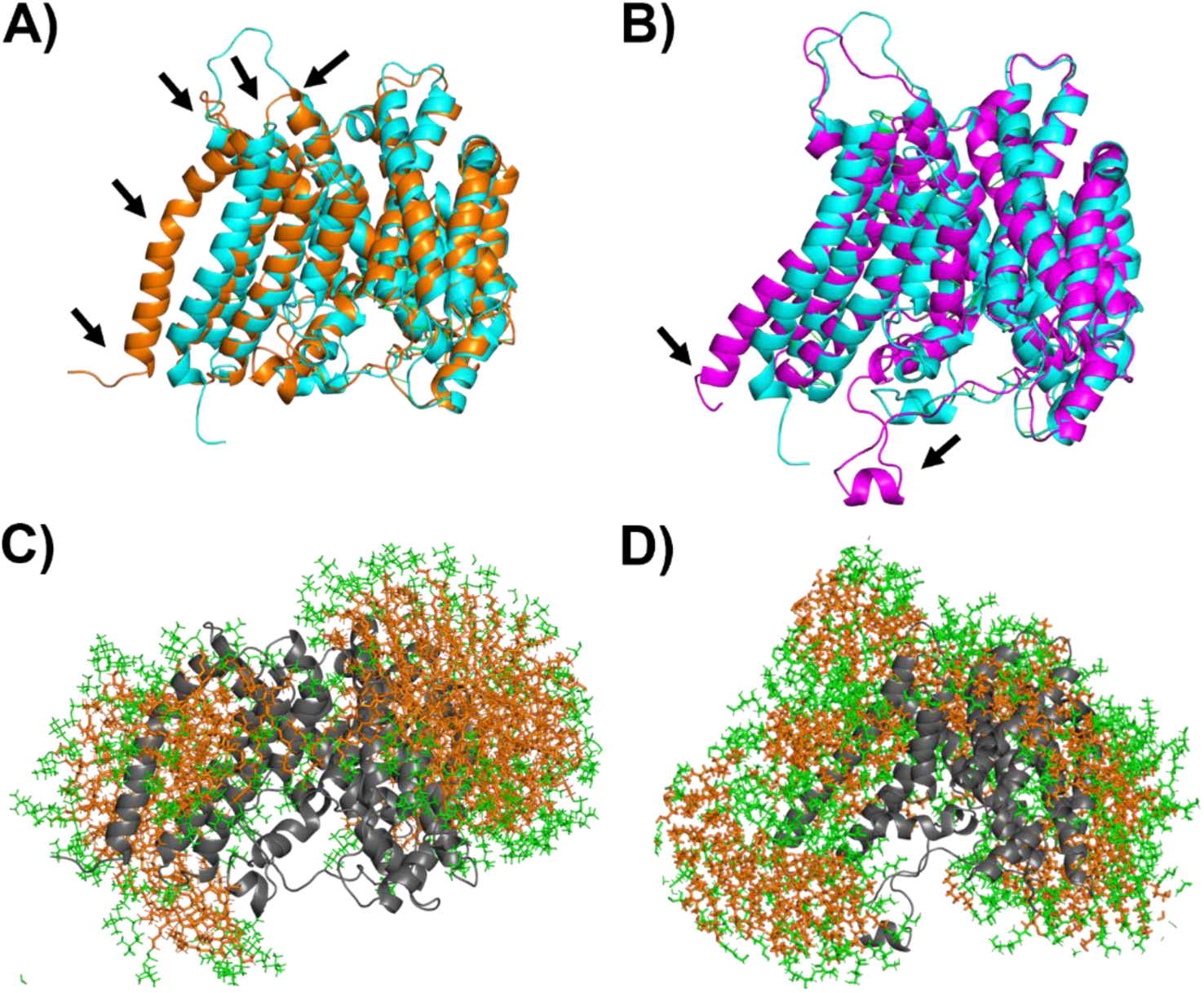
Comparison of hGOAT stabilization and detergent interactions. **A and B)** Structural alignment of hGOAT structure (cyan) with hGOAT solubilized by (A) FOS-16 (orange) and (B) MEGA-9 (purple). Black arrows indicate regions of highest deviation between the structures as reflected by RMSF. **C and D)** Interactions of hGOAT (gray, cartoon) with hydrophilic head groups (green, sticks) and hydrophobic tail groups (orange, sticks) when solubilized by (A) FOS-16 and (B) MEGA-9 exhibit distinct patterns with more head group interactions with the non-ionic MEGA-9 polyol than the FOS-16 phosphocholine zwitterion.

### Construction of a hGOAT-eGFP fusion protein for monitoring detergent solubilization by in-gel fluorescence

Recent studies using attachment of fluorescent proteins to mammalian integral membrane proteins have demonstrated this approach facilitates detergent solubilization studies by enabling protein detection by in-gel fluorescence and fluorescence size exclusion chromatography (FSEC).^40, 44^ We developed a hGOAT-eGFP construct with a C-terminal His_10_ tag to allow fluorescence-based detection and IMAC-based protein purification following solubilization (Figure 4A). This construct expresses well in our baculovirus system,^44^ with in-gel fluorescence detected at an apparent molecular weight of ∼55 kDa and anti-MBOAT4 western blotting revealing two bands at ∼70 kDa and ∼55 kDa (Figure 4B). This banding is consistent with a partially denatured protein with intact fluorescent eGFP detected in the lower band and the fully denatured protein running higher without eGFP fluorescence.^48–51^

**Figure 4.**
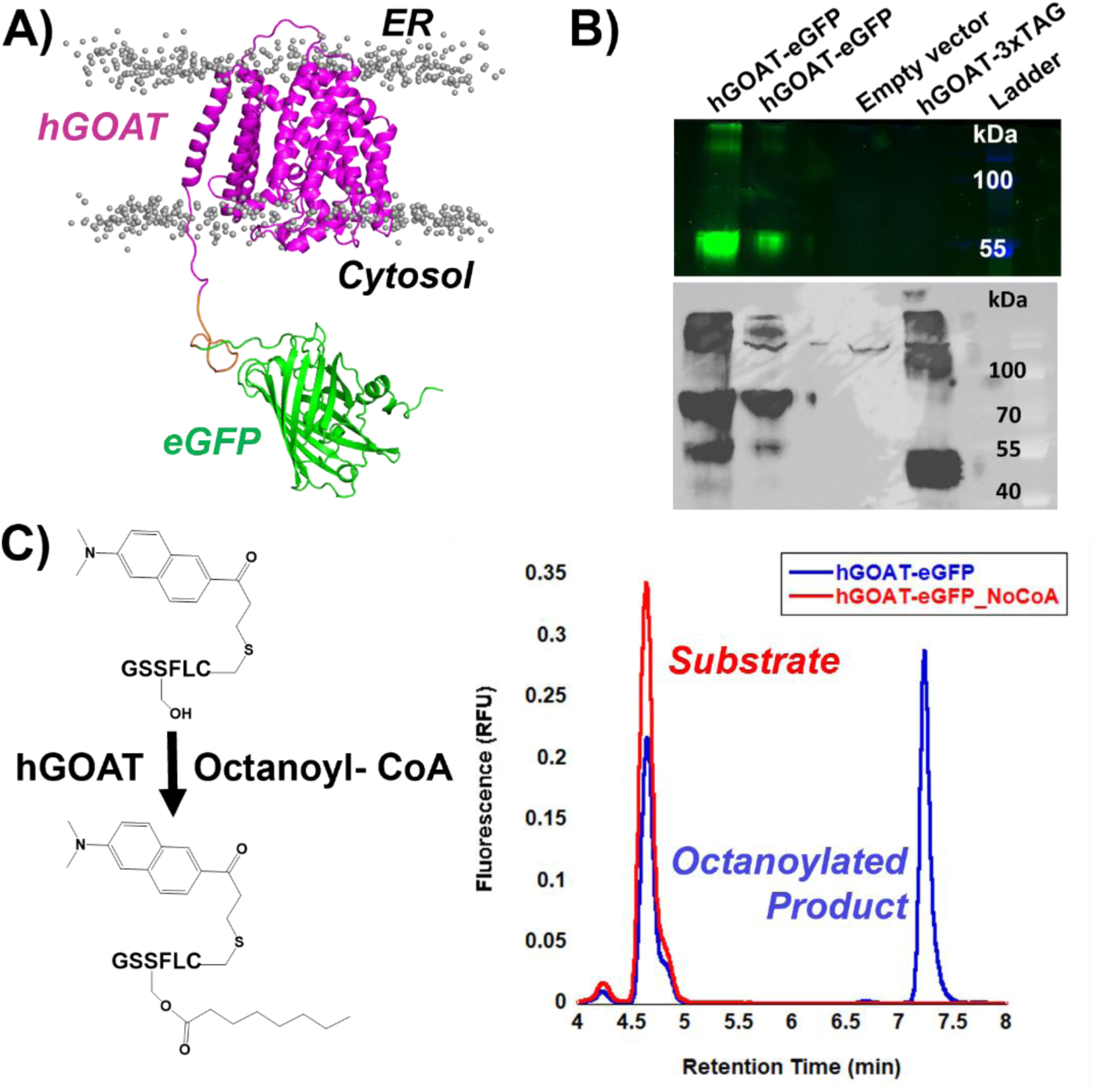
Expression and activity validation of a hGOAT-EGFP construct. To allow fluorescence-based detection, an eGFP tag was appended to the C-terminus of human GOAT (hGOAT). Addition of this tag preserved expression and activity of GOAT. **A)** Structural model of the hGOAT-eGFP fusion protein embedded in a phospholipid bilayer. Lipid headgroups are shown as grey sphere, and lipid tails are omitted for clarity. **B)** In-gel fluorescence detection of hGOAT-eGFP. The gel was imaged with the Alexa488 filter (samples) and Coomassie Blue (ladder) and these two filter images were overlayed. The presence of the 55kDa band for hGOAT-eGFP is consistent with a partially denatured protein maintaining the eGFP fold, as described in the text. No fluorescence was observed for the empty vector control and our previously published hGOAT-3xTAG construct. **C)** Anti-MBOAT4 immunoblot detects both a 70kDa and 55kDa band corresponding to the fully denatured hGOAT-eGFP at 70kDa and the GOAT denatured by eGFP still fluorescent band at 55kDa. Our previously published GOAT construct hGOAT-3xTAG (57kDa; running size 45kDa) serves as a positive control for the antibody.^44^ **D)** hGOAT-eGFP construct catalyzed ghrelin acylation. In the presence of enzyme and octanoyl-CoA, the substrate peptide is acylated to yield the more hydrophobic octanoylated product as monitored by reverse phase HPLC with fluorescence detection.

The ghrelin acylation activity of the hGOAT-eGFP fusion protein was determined using a fluorescently labeled ghrelin mimetic peptide in our established HPLC assay (Figure 4C).^40, 43–44^ The fusion protein exhibits robust acylation activity comparable to the non-eGFP tagged construct commonly used in our studies, demonstrating this construct is compatible with determining both protein solubilization and enzyme activity in detergent screens.

### Detergent solubilization screen of hGOAT-eGFP

Solubilization trials with hGOAT-eGFP explored the same eight detergents analyzed in the computational studies, with detergents at 1 % (w/v) and 4% (w/v) in each trial. Following incubation with each detergent with rotation at 4 °C and separation of solubilized and membrane-resident proteins by ultracentrifugation, soluble proteins in the supernatant and insoluble proteins in the pellet were analyzed by SDS-PAGE in gel fluorescence. Each trial included a buffer-only negative control for hGOAT-eGFP solubilization, and each gel contained a lane with untreated hGOAT-eGFP membrane fraction to provide a positive control for hGOAT-eGFP in-gel fluorescence and western blotting (Figure 5). FOS-16 effectively solubilized hGOAT-eGFP as expected based on previous studies of the mouse GOAT isoform (Figure 5A),^32^ and FOS-12 with a shorter alkyl chain similarly solubilized hGOAT-eGFP to a large extent. Partial solubilization was observed with CHAPS, LMNG, GDN, and DDM, whereas BOG and MEGA-9 exhibited minimal ability to solubilize hGOAT-eGFP (Figure 5B). The solubilized supernatant fraction from each solubilization trial were assayed for ghrelin acylation activity, with the unsolubilized pellet from the buffer-only control providing a positive control for enzymatic activity. Of the detergent supernatant fractions, only the MEGA-9 supernatant exhibited acylation activity with a ghrelin-mimetic peptide (Figure 6). This suggests that a small concentration of MEGA-9 solubilized hGOAT-eGFP retains sufficient native fold to support ghrelin binding and acylation.

**Figure 5.**
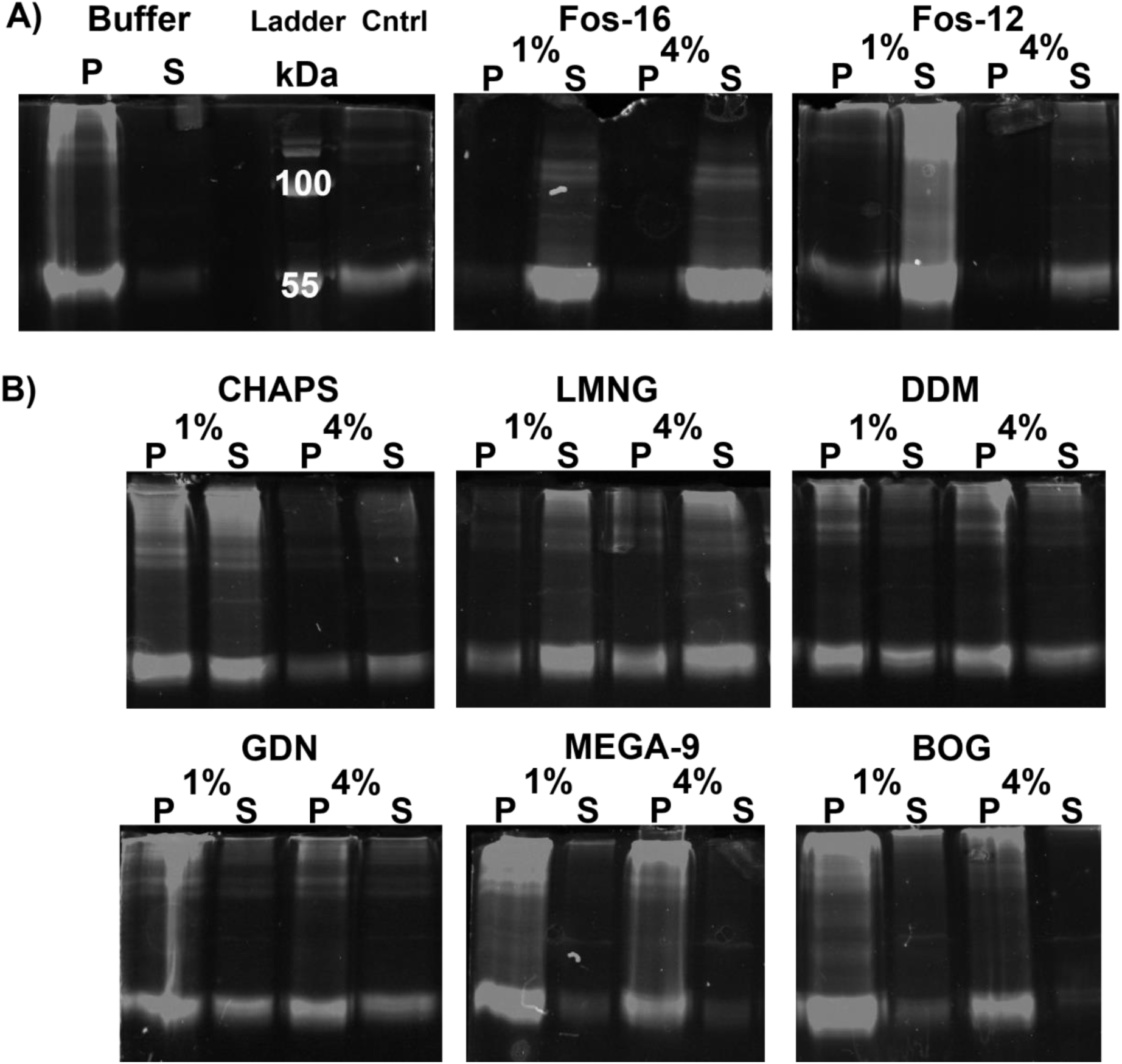
hGOAT-eGFP solubilization monitored by in-gel fluorescence. A) Buffer sample negative control and untreated hGOAT-eGFP negative control provide size standard for fluorescent hGOAT-eGFP, with FOS-16 and FOS-12 exhibiting efficient solubilization of hGOAT-eGFP as shown by majority of fluorescence in the supernatant/soluble protein fraction. B) hGOAT-eGFP solubilization trials exhibited partial solubilization with CHAPS, DMM, and GDN, more effective solubilization with LMNG, and little to no solubilization with MEGA-9 and BOG.

**Figure 6.**
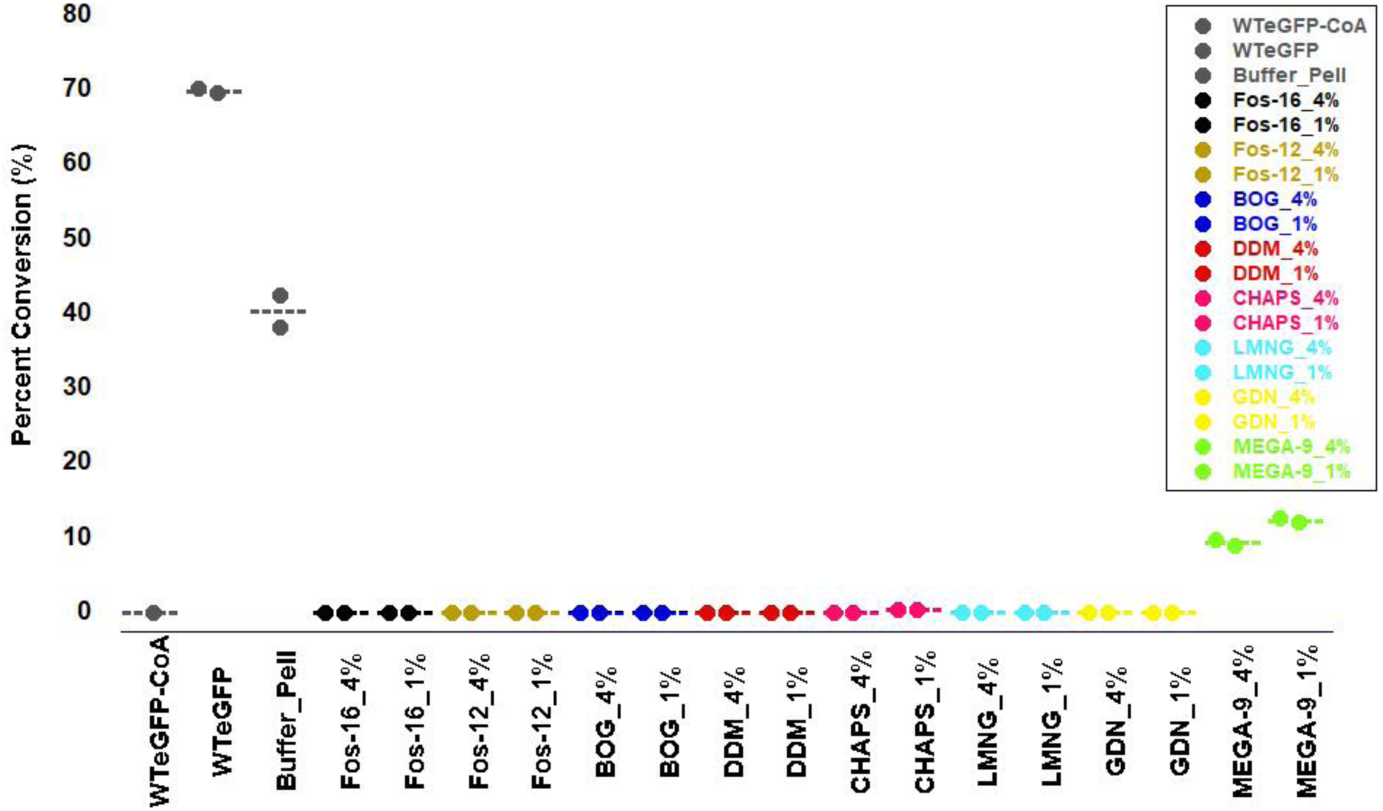
MEGA-9 maintains octanoylation activity in its hGOAT-eGFP supernatant fraction. Each hGOAT-eGFP solubilization supernatant fraction was assessed for ghrelin octanoylation activity. Reaction lacking the acyl donor served as a negative control, with untreated WT hGOAT-eGFP and the pellet from the buffer-treated hGOAT-eGFP serving as positive controls. Only the supernatant fraction from the MEGA-9 solubilizations exhibited significant activity with ∼20% conversion of substrate to octanoylated product. Activity screening reactions were performed in duplicate and analyzed as described in Experimental Methods.

Based on the solubilization and enzyme activity results from this initial screen, we expanded our analysis of several detergent families to determine if a related detergent could provide additional solubilization of hGOAT-eGFP or support hGOAT-eGFP acylation activity (Supporting Figures S1 and S2). Based on the acylation activity exhibited by the MEGA-9 trial supernatant, we examined the related detergents HEGA-11 and MEGA-10 to determine if they could more efficiently solubilize hGOAT-eGFP while maintaining enzyme activity. Unfortunately, neither of these detergents were more effective than MEGA-9 in solubilizing hGOAT-eGFP. Similar expansion from LMNG to the related detergents DMNG and OGNG did not result in increased solubilization or enzyme activity. Finally, trials with a series of FOS family detergents with decreasing alkyl chain lengths showed that a minimum alkyl chain length of ten carbons is required for complete solubilization. The shorter chain detergents FOS-9 and FOS-8 did not support efficient solubilization but exhibited detectible hGOAT acylation activity in their solubilized supernatant.

### Analysis of solubilized hGOAT-eGFP polydispersity by FSEC

Optimization of the detergent:total protein ratio with our top four performing detergents (GDN, LMNG, DDM, and CHAPS) resulted in efficient solubilization of hGOAT-eGFP in each case comparable to that observed with FOS-16 (Figure 7A). We analyzed the solubilized hGOAT-eGFP samples by fluorescence size exclusion chromatography (FSEC) which provides information regarding the monomeric, oligomeric, or aggregated state of solubilized membrane proteins.^35, 52^ Under our separation conditions, a fluorescence peak at a retention volume of ∼8 mL indicates a protein aggregate eluting in the column void volume. hGOAT-eGFP solubilized in FOS-16 elutes as a single peak at a retention volume of 15.3 mL, consistent with previous reports of FOS-16 solubilization resulting in monomeric GOAT.^34^ CHAPS solubilized hGOAT-eGFP exhibited significant aggregation as reflected in a peak at ∼ 8mL without major peaks at larger retention volumes consistent with discrete monomers or oligomers. In contrast, hGOAT-eGFP solubilized in the other three detergents exhibited FSEC profiles consistent with a mixture of monomeric and dimeric/oligomeric species (Figure 7B). The DDM-solubilized sample exhibited a wide peak with two maxima at 17.6 mL and 18.1 mL. hGOAT-eGFP solubilized in GDN and LMNG displays more discrete monomer/oligomer distributions with two peaks in GDN (16.3 and 17.6 mL) and LMNG (15.3 and 18.2 mL). We note the lower integrated intensity of the DDM, GDN, and LMNG FSEC peaks compared to FOS-16 indicates further effort is needed to maximize the yield of solubilized hGOAT with these detergents. These promising findings provide the foundation for further studies towards experimental structural characterization of GOAT.

**Figure 7.**
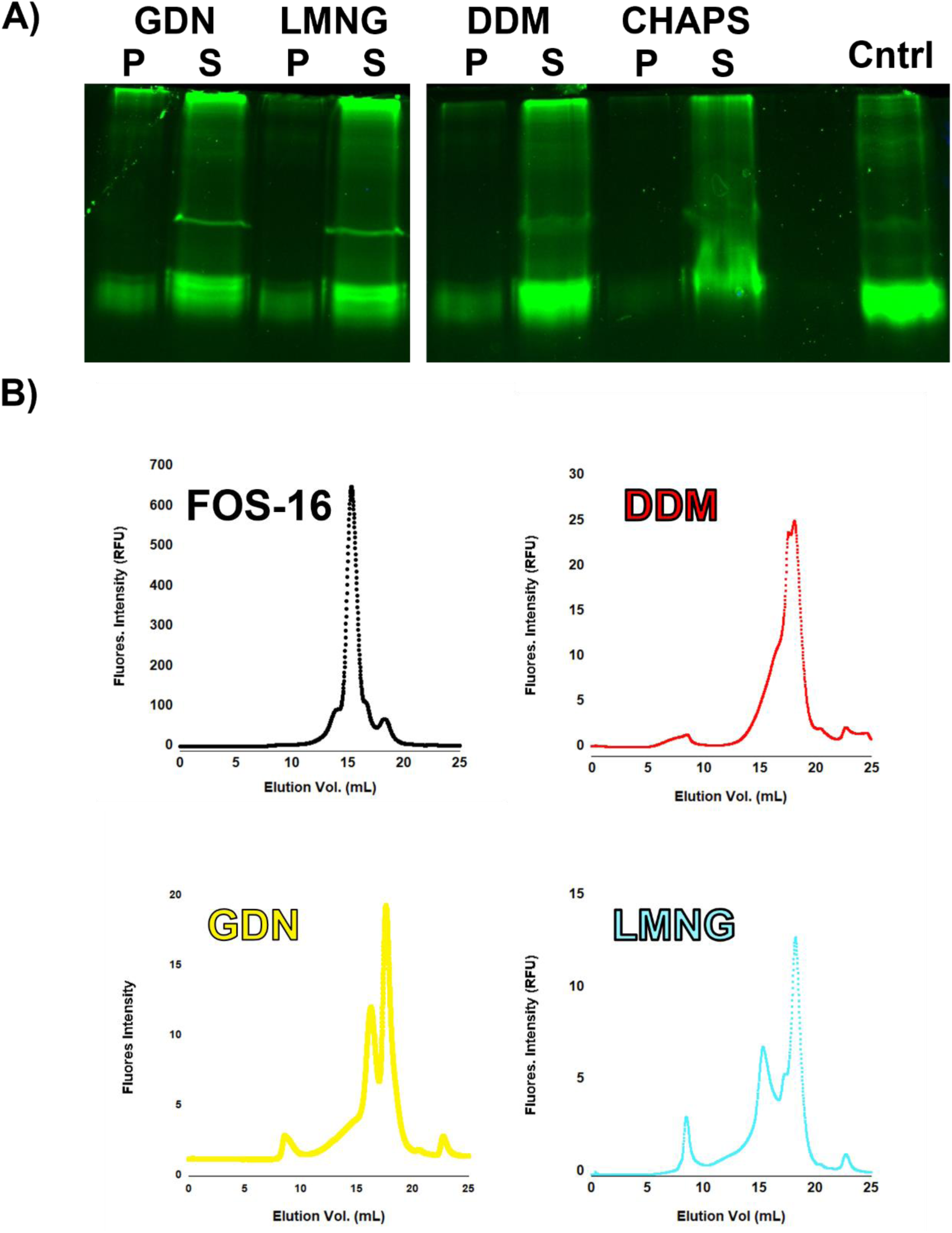
FSEC analysis indicates polydispersity for hGOAT-eGFP solubilized in DDM, GDN, and LMNG. A) Optimized conditions increase hGOAT-eGFP solubilization in GDN, LMNG, DDM, and CHAPS B) FSEC analysis indicates a single peak for FOS-16 solubilized hGOAT-eGFP, while hGOAT-eGFP in DDM, GDN, and LMNG exhibit multiple peaks indicating enzyme-detergent complex polydispersity. Representative chromatograms reflect solubilizations run in triplicate on different days.

## Discussion

Here we describe our approach to developing a computational framework informing integral membrane protein detergent solubilization. We analyzed the two major requirements for this process – detergent solubilization/invasion of the phospholipid bilayer and detergent stabilization of the membrane protein folded tertiary structure. We found that the rank order of membrane invasion ability did not correspond to our biochemical data for detergent solubilization of hGOAT-eGFP. In contrast, the analysis of protein stabilization by the various detergents more closely matched our experimental findings with the detergents CHAPS, LMNG, and GDN amongst the best performing detergents in both analyses. Unfortunately, these same detergents support very little GOAT activity in our ghrelin acylation assay. This lack of activity could reflect insufficient yield of solubilized hGOAT-eGFP or detergent inhibition of hGOAT enzymatic activity. In addition, we observe hGOAT acylation activity in the supernatant of the MEGA-9 solubilization trial, whereas very little solubilized hGOAT-eGFP was observed by in-gel fluorescence (Figures 5-6). This observation supports the second explanation that LMNG, GDN, and CHAPS inhibit hGOAT activity while exhibiting superior protein solubilization. This inhibition could arise from direct interference with substrate binding, disruption of active site structure, larger-scale disruption of enzyme structure and/or dynamics required for catalysis, or a superposition of these three inhibition mechanisms. Future studies of ghrelin acylation by hGOAT and hGOAT inhibition in the context of purified enzymes may be aided by transfer into MEGA-9 and similar detergents following purification. Such requirements for detergent exchange to support integral membrane enzyme activity are commonly reported, as in the case of other MBOAT family enzymes.^23, 35, 38, 53–57^

In the case of the fos choline detergents, expansion upon the initial detergent screening provides further insight into the specific characteristics supporting both hGOAT solubilization and enzymatic activity. As the alkyl chain was shortened from sixteen to eight carbons, a reduction in hGOAT-eGFP solubilization was accompanied by an increase in ghrelin acylation activity of the solubilized enzyme most notable with FOS-9 and FOS-8. This behavior likely reflects a trade-off of detergent solubilization ability and perturbation of enzyme structure upon solubilization leading to a loss of catalytic activity. We did not observe similar changes in the MEGA/HEGA and neopentyl glycol detergent families, wherein we explored changes in both hydrophobic alkyl chains and the hydrophilic amine and glycoside head groups.

Our top-performing detergents GDN, LMNG, and DDM exhibit multiple peaks depicting oligomeric behavior and some aggregation of hGOAT-eGFP when solubilized hGOAT-eGFP was analyzed by size exclusion chromatography (Figure 7b). This polydispersity reinforces the need to identify improved detergent systems that maintain hGOAT activity and solubility in a monodisperse protein-detergent complex. Based on structural studies of the other protein modifying MBOAT family members, PORCN and HHAT,^35, 37–38^ GOAT likely exists as a monomer. Analysis of potential hGOAT dimer formation using the PANEL computational approach similarly supports GOAT existing a monomer (data not shown).^58^The size of monomeric GOAT alone will be insufficient to support structural studies by cryo-electron microscopy without formation of a larger complex with binding partners such as GOAT-targeted antibodies or other specific binding partners as has been used with related integral membrane proteins.^23^

Determining which detergent to use for membrane protein solubilization, purification, and structural analysis can require a significant amount of time and laboratory resources. Most mammalian membrane proteins are difficult to express in large quantities for detection, let alone in purification quantities. This adds tremendous difficultly when solubilizing mammalian membrane proteins. We have developed a computational approach to “whittle down” the detergent choices leading to more efficient biochemical screening. This approach requires a computational model of the protein interest. Fortunately, this requirement is less of an issue with the emergence of AlphaFold and related approaches which can provide a reasonable starting point for structural modeling and analysis.^20^ We believe our approach presents a new way to quickly look for detergents compatible with a specific protein of interest.

## Methods

### General methods

Data plotting was carried out with Kaleidagraph (Synergy Software, Reading, PA, USA). Antibiotics and LB Media for DNA and bacmid propagation were purchased from BioBasic and ThermoFisher.

### Computational methods

To study the invasion of eight types of detergents on phospholipid bilayer, a bilayer of DOPC (dioleoylphosphatidylcholine) and DPPC (dipalmitoylphosphatidylcholine) was built with a ratio of 1:1 from CHARMM-GUI Membrane Builder.^46–47^ The number of DOPC and DPPC molecules in the bilayer is 80. The atomistic structures of all eight detergent molecules in this work were obtained from CHARMM-GUI Ligand Reader & Modeler.^59–60^ For each detergent, 240 molecules were inserted into a cubic simulation box (12 nm length) with the bilayer, and thus the number ratio of detergent to lipid is 3:1. To study the stability of GOAT structure with detergents, another eight simulation systems with the box length of 15 nm were built, in which each contained one GOAT protein and 490 detergent molecules. Details regarding generation of the computational model for human GOAT structure were discussed in our previous work.^17^

These systems were subjected to a series of energy minimization and equilibration steps with the input files generated from CHARMM-GUI solution builder.^59, 61–62^ The CHARMM36m force field parameters were used for GOAT protein, lipids, detergents, salt (0.15M NaCl), and explicit TIP3P water.^63^ The atomistic molecular dynamics simulations were carried out using the GROMACS version 2019.^964^ Each system was energy minimized, followed by equilibration in isothermal-isochoric (*NVT*) and isothermal-isobaric (*NPT*) for 1 ns each, and production MD run under *NPT* conditions for 500 ns in studying the invasion process of detergents on phospholipid bilayer. The production MD run was 200 ns in studying the stability of GOAT structure. The heavy atoms of the GOAT proteins were restrained during *NVT* and *NPT* equilibration. All restraints were removed during the production MD. The temperature was maintained 298 K using the v-rescale thermostat with τ_t_ = 1.0 ps.^65^ In the pre-production *NPT* run, isotropic pressure of 1 bar was maintained using Berendsen barostat^11^ with τ_p_ = 5.0 ps and compressibility of 4.5×10^−5^ bar^−1^.^66^ In the production MD, we used the Parrinello-Rahman barostat with τ_p_ = 5.0 ps and compressibility of 4.5×10^−5^ bar^−1^.^67^ Three-dimensional periodic boundary conditions (PBC) were applied to each system. A 2 fs time step was used, and the nonbonded interaction neighbor list was updated every 20 steps. A 1.2 nm cutoff was used for the electrostatic and van der Waals interactions. The long-range electrostatic interactions were calculated using the Particle-Mesh Ewald (PME) method after a 1.2 nm cutoff. The bonds involving hydrogen atoms were constrained using the linear constraint solver (LINCS) algorithm.

Molecular visualization and images were rendered using PyMol and VMD software suites.^68–69^ Data analysis and plotting were performed using in-house Python scripts based on publicly hosted python packages, such as matplotlib, scipy, and MDAnalysis.^70^

### hGOAT-eGFP cloning and baculoviral expression

Primers were ordered from Integrated DNA Technologies (IDT) to insert a *Xho*I restriction endonuclease at the N-terminus of eGFP in pEGBACMAM_hGOAT-eGFP vector,^52^ (JH_pEGBACMAM_FL_Xho1_For (5’-CAGTCTCGAGGTGAGCAAGGGCGAGGAGCTG-3’) and JH_pEGBACMAM_FL_Rev (5’-CAGAGGTTGATTAAGCTTGTCGAGACTGCA-3’)) and dissolved in ultrapure water. Primer concentrations were determined by absorbance at 260 nm. PCR reactions (50 µL total volume) contained 5x Standard Buffer, 10 mM dNTPs, 10 µM of forward and reverse primers, 10 ng/µL of template DNA, and 1 µL TaqOne polymerase. PCR reactions ran for 32 cycles, with an initial denaturation cycle (95°C, 1 min), 30 cycles of denaturation (95°C, 30 sec); annealing (56°C, 60 sec) and extension (68°C, 2 min) and one final extension cycle (68°C, 5 min). The PCR reaction mixture was analyzed by agarose gel electrophoresis (0.8% agarose, 1X TAE buffer) and imaged with a Bio-Rad Molecular Imager ChemiDoc™ XRS+ camera with Image Lab 4.1 software. PCR reactions were purified by EZ-10 Spin Column DNA PCR Purification Kit (Bio Basic Inc. #BS664-120712) with elution by 30 µL of ultrapure water instead of 50 µL elution buffer.

The PCR product encoding eGFP-hGOAT was cloned into the pFastBacDual vector using the *Xho*I and *Xba*I restriction sites. Insertion of the hGOAT-eGFP gene was verified by DNA sequencing (Genewiz). Preparation of eGFP-hGOAT baculovirus was performed using the Bac-2-Bac protocol (Invitrogen), with the presence of the hGOAT-eGFP insert verified by colony PCR amplification and PCR of the purified baculovirus as previously described.^44^ hGOAT-eGFP expression and membrane fraction enrichment was performed as previously described.^44^

### hGOAT-eGFP acylation activity assay

Assays were performed and analyzed by reverse-phase HPLC as previously described.^44^ Octanoyl coenzyme A (octanoyl-CoA, free acid, Advent Bio) were solubilized to 5 mM in 10 mM TrisHCl (pH 7.0), aliquoted into low-adhesion microcentrifuge tubes, and stored at −80 °C. Unlabeled GSSFLC_NH2_ peptide was synthesized by Sigma-Genosys (The Woodlands, TX), solubilized in 1:1 acetonitrile:H_2_O, and stored at -80 °C. Acrylodan (Anaspec) for peptide substrate labeling was solubilized in acetonitrile with the stock concentration determined by absorbance at 393 nm in methanol (ε_393_ = 18,483 M^−1^ cm^−1^ , per the manufacturer’s data sheet). GSSFLC_NH2_ peptide concentrations were determined by reaction of the cysteine thiol with 5,5’-dithiobis(2-nitrobenzoic acid) and absorbance at 412 nm, using ε412 = 14,150 M^−1^ cm^−1^ .^71^ Peptide substrate fluorescent labeling was performed using published protocols.^44^

Following acylation reactions, the fluorescent peptide substrate and octanoylated product were detected by fluorescence (λ_ex_ 360 nm, λ_em_ 485 nm). Chromatogram analysis and peak integration was performed using Chemstation for LC (Agilent Technologies). Product conversion was calculated by dividing the integrated fluorescence for the product peak by the total integrated peptide fluorescence (substrate and product) in each run.

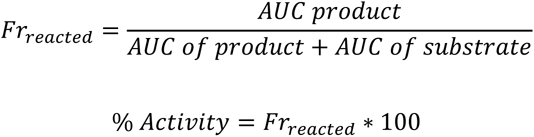

### Polyacrylamide gel electrophoresis and in-gel fluorescence of hGOAT-eGFP

For analysis by gel electrophoresis, 20-50 µg of membrane protein fraction (concentration determined by Bradford 1X Dye reagent (Bio-Rad)) in a total volume of 15-30 uL 1x gel loading sample buffer (250 mM Tris HCl pH 6.8, 8% SDS (w/v), 40% glycerol (v/v), 5% BME (v/v), 0.4 % bromophenol blue (w/v)) is heated to 50.2 °C for 5 min. Samples were loaded onto a 10% polyacrylamide gel with 1x Tris-glycine buffer (25 mM Tris Base, 150mM Glycine, 1% SDS, pH 8.1-8.8) and electrophoresed at 160V. In detergent solubilization experiments, untreated hGOAT-eGFP membrane fraction is loaded as a control for hGOAT-eGFP size.

All expression analysis and detergent solubilization gels with hGOAT-eGFP are analyzed by in-gel fluorescence detection. Gels were imaged with a Bio-Rad Molecular Imager ChemiDoc™ MP Imaging System using Filter Chemiluminescence and Image Lab 4.1 software. To visualize hGOAT-eGFP filters Pro-Q Emerald 488 or Alexa488 filter were used for the fluorescently tagged proteins and Coomassie Blue* filter to visualize the ladder in the gel. These two images were then merged for analysis.

### hGOAT-eGFP immunoblotting

For Western blotting, following gel electrophoresis proteins were transferred from the gel to a polyvinylidene fluoride (PVDF) membrane using a Trans-Blot Turbo Transfer System (BioRad, Trans-Blot turbo RTA transfer kit) using the Bio-Rad standard protocol (25 V,30 min). The PVDF membrane was blocked with EveryBlot Blocking Buffer (BioRad;#12010020) for 30min-2hr. The membrane was incubated in 1:1000 1° MBOAT4 Polyclonal antibody (Cayman Chemical #18614) and 10 mL of EveryBlot Blocking Buffer (BioRad;#12010020), overnight at 4°C or 1-2hr at RT, rocking. Post incubation, the membrane is washed in 1xTBST six times for five minutes a piece. Secondary antibodies are diluted in 10ml EveryBlot Blocking Buffer (BioRad;#12010020) with 1:2500 goat anti-rabbit-HRP (Invitrogen; ref#65-6120) for MBOAT4 primary. Once secondary is added, these are rocked for 1hr at RT. Post incubation, the membrane is washed in 1xTBST six times for five minutes a piece. The PVDF membrane is exposed to chemiluminescent substrates in a 1:1 ratio (ThermoScientific #34577) for 5 min. Gels were imaged with a Bio-Rad Molecular Imager ChemiDoc™ MP Imaging System; Filter Chemiluminescence for antibody bound constructs and the Colorometic filter to image the ladder. These two images are then merged and analyzed using Image Lab 4.1 software.

### Detergent solubilization screening trials

Detergents were ordered as part of a solubilization screening kit (Popular Detergent Kit; 850561P-1EA, Avanti Polar Lipids) or individually for Fos-Choline-16 (F316, Anatrace) and Lauryl Maltose Neopentyl Glycol (NG310, Anatrace). Detergents were solubilized in solubilization buffer: 50mM HEPES pH 7, 500 mM NaCl, and 10% glycerol.^72^

For solublization trials, hGOAT-eGFP membrane fraction was thawed on ice. Detergents were added in a 1:1 (vol./vol.) ratio to yield a 1% or 4% detergent solution concentration. Samples were rotated in ultracentrifuge grade 1.5 mL Eppendorf tubes for 2 hrs at 4°C. Samples were then ultracentrifuged at 38,000 x *g* for 30min at 4°C. Following centrifugation, the supernatant was removed to a new microcentrifuge tube and the pellet was gently resuspended in 100 µL of solubilization buffer. For in-gel fluorescence analysis, supernatant or pellet resuspension (30 µL) was combined with 3x sample loading buffer and heated at 50°C for 5 min before loading onto a 10% SDS polyacrylamide gel. Gels were imaged and analyzed as described above.

For subsequent optimization experiments, solubilizations with DDM, GDN, and LMNG were performed with a detergent:protein mass ration of 20:1 at a detergent concentration of 4% (w/v). All other steps were performed identically to the protocol above.

### Fluorescence size exclusion chromatography (FSEC) analysis of detergent solubilized hGOAT-eGFP

2 mg/mL of total protein (DC Bradford; cat. 5000111) including detergent solubilized hGOAT-eGFP (∼300 µL) was filtered through a 0.2 µm PES filter (Cytiva, Whatman UNIFLO 13mm cat#99142502). 100 µl of the filtered sample was diluted with 100 µL of the detergent-specific FSEC buffer (40 mM HEPES pH 7.5, 150 mM NaCl, 5% glycerol, 0.08-0.01% detergent, 1 mM BME), and 100 µL of the diluted sample was injected into the FPLC. For hGOAT-eGFP solubilized in FOS-16, 100 µL of undiluted filtered sample was injected. All FSEC experiments were performed on a Cytiva AKTA pure 25 chromatography system equipped with a Superose 6 increase 10/300 GL column (ca. t#29091596, Cytiva), integrated UV-Vis detector, and an in-line RF-20A Shimadzu fluorescence detector. FSEC analysis was run at 0.30 mL/min flow rate with detergent-specific FSEC buffers (40 mM HEPES pH 7.5, 150 mM NaCl, 5% glycerol, 0.08-0.01% detergent, 1 mM BME). Fluorescence (λ_ex_ 485 nm, λ_em_ 512 nm) was monitored for eGFP tagged constructs, with detector response values exported using Unicorn software (Cytiva) and replotted using Kaleidagraph (Synergy Software, Reading, PA, USA).

## Data and Software Availability Statement

Coordinates for the hGOAT structural model are available upon request from the corresponding authors. Molecular dynamics simulations were performed using GROMACS (https://www.gromacs.org). The topology pdb files for all detergents and the gromacs mdp files for membrane invasion and protein stability are available at https://github.com/NangiaLab/GOAT_detergents

## Acknowledgements

The authors gratefully acknowledge members of the Nangia and Hougland research groups for discussion and comments on the manuscript. This study was supported by the National Institutes of Health (grant GM134102) and the National Science Foundation (CAREER CBET-1453312, NSF DMR-BMAT-2105193, NSF-DMR-XC-1757749 and 2049793, NSF-MCB-2221796). Computational resources were provided by the Information and Technology Services at Syracuse University and the Extreme Science and Engineering Discovery Environment (XSEDE), supported by National Science Foundation grant number ACI-1053575. Anton 2 computer time was provided by the Pittsburgh Supercomputing Center (PSC) through Grant R01GM116961 from the National Institutes of Health.

## Supporting Information

hGOAT-eGFP solubilization trials using detergent family expansions from initial hit screens; hGOAT activity analysis of solubilization trials using detergent family expansions from initial hit screens (PDF)

## For Table of Contents Only

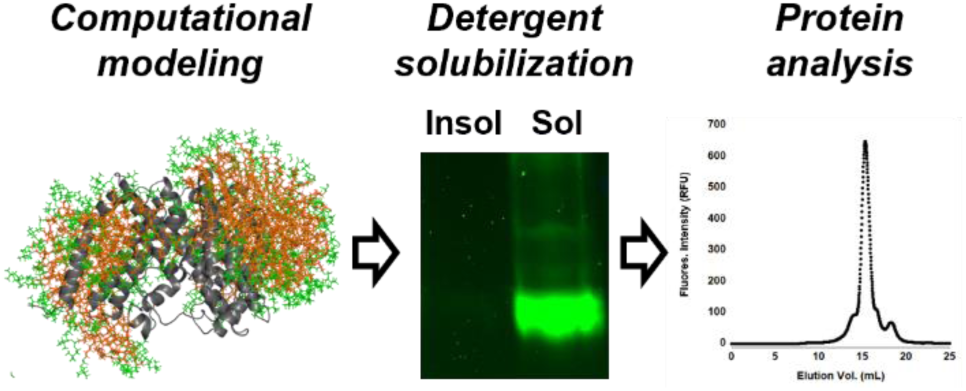

