## Supporting Information for "A combined computational-biochemical approach offers an accelerated path to membrane protein solubilization"

**Supporting Figure S1.** MEGA-9, LMNG, and Fos-choline family expansions do not improve solubilization.

**Supporting Figure S2.** MEGA-9, LMNG, and Fos-choline family expansions showed minimal improvement in hGOAT-eGFP activity.

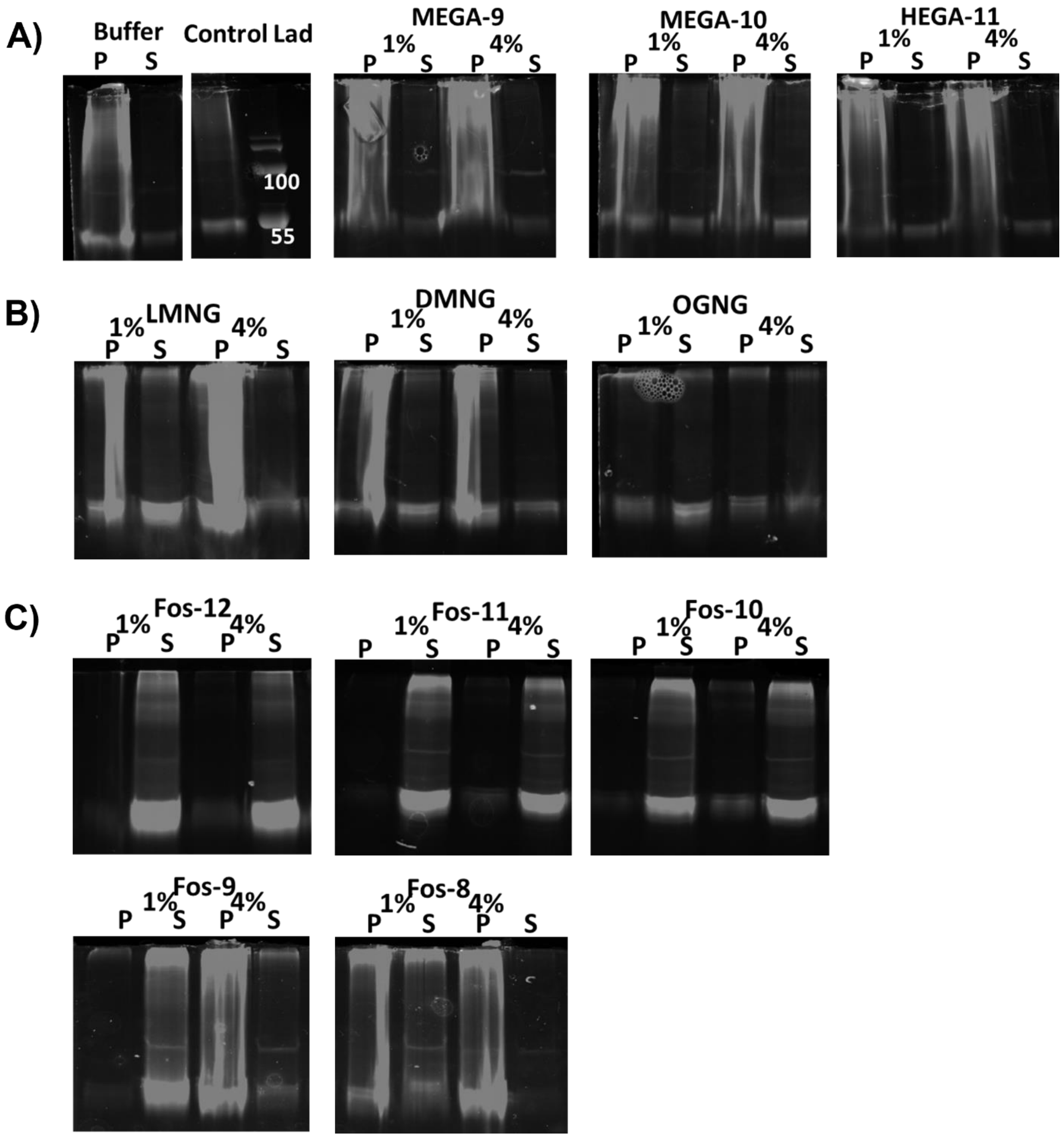


**Supporting Figure S1. MEGA-9, LMNG, and Fos-choline family expansions do not improve solubilization.** A solubilization screening was completed with the expanded MEGA-9, LMNG, and Fos-choline families. Buffer control is identical to that performed for the detergent solubilizations in the main text. A) MEGA-9 had comparable solubilization to the detergent screen and MEGA-10 did not improve the solubilization. HEGA-11 depicted does not show solubilization in either detergent percentage. B) DMNG showed the majority of the solubilization in the pellet fraction with some hGOAT-eGFP in the supernatant fractions. OGNG shows partial solubilization, but is not improved from the solubilization from LMNG. C) Fos-12 repeated its solubilization pattern from the initial screen, and the foc-choline detergents with shorter alkyl chain lengths show a gradual decrease in hGOAT-eGFP solubilization.

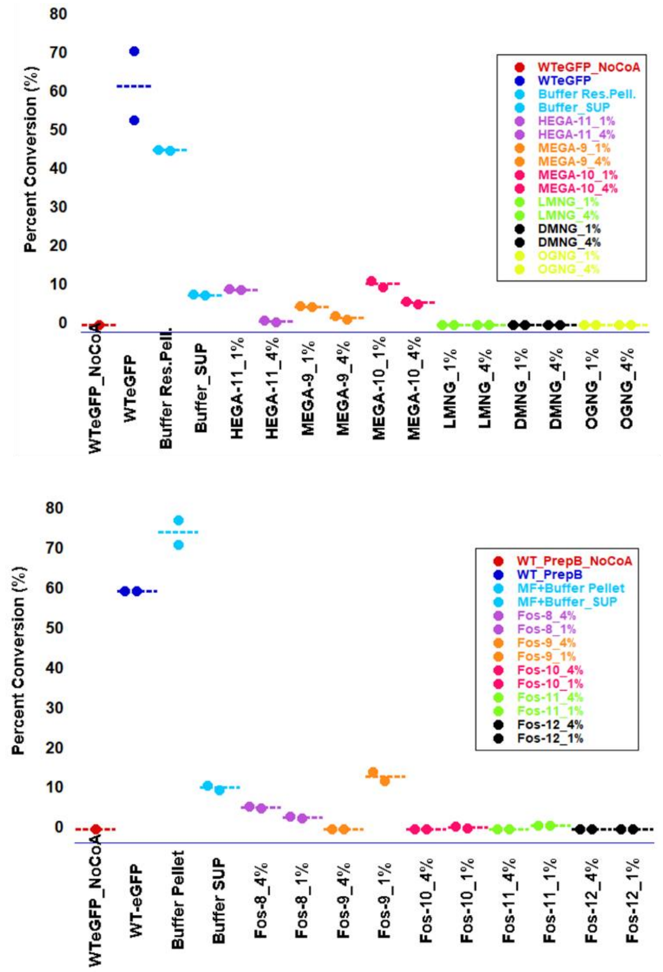


**Supporting Figure S2. MEGA-9, LMNG, and Fos-choline family expansions showed minimal improvement in hGOAT-eGFP activity.** Supernatant fractions of the solubilization reactions tested for ghrelin acylation activity as described in the main text, with negative and positive controls. hGOAT-eGFP solubilized by HEGA-11, MEGA-10, and MEGA-9 all show similar reactivity to the sample without detergent whereas LMNG, DMNG, and OGNG have inhibited hGOAT-eGFP activity. Amongst the fos-choline detergents, only FOS-9 solubilized hGOAT-eGFP exhibited appreciable acylation activity. Activity screening reactions were performed in duplicate and analyzed as described in Experimental Methods.
